# The evidence of trace amounts of methane of uncertain origin released from cerambycid larvae of *Stictoleptura rubra*

**DOI:** 10.1101/2023.03.31.535042

**Authors:** Aleksandra Walczyńska, Kevin Purdy, Jarosław Nęcki

**Affiliations:** Institute of Environmental Sciences, Jagiellonian University, Gronostajowa 7, 30-387, Krakow, Poland; School of Life Sciences, University of Warwick, Gibbet Hill Campus, Coventry, CV4 7AL, UK; Faculty of Physics and Applied Computer Science, AGH - University of Science and Technology, Mickiewicza 30, 30-059 Krakow, Poland

**Keywords:** Archaea, Cerambycidae, methanogenesis, symbiosis, wood-feeding

## Abstract

Insects release methane as a result of their symbiosis with methanogenic microorganisms. This phenomenon has been well studied in termites but is still poorly known in other insects, among which the most likely methane producers are wood-feeders. We applied two methodological approaches to examine whether the wood-feeding larvae of *Stictoleptura rubra* (Cerambycidae, Coleoptera) release methane. By means of the precise gas chromatography we detected a methane release at the rate of 0.02-0.05 nmol/g×hr. We also investigated the gut Archaea assemblage by 16S rRNA gene sequencing and T-RFLP analysis. Halophilic archaea but no known methanogens were detected. Combined with our chromatographic studies showing small but significant amount of methane release, one possible explanation is that the detected archaeons are methanogenic although this is unlikely. Our results offer the first evidence of methane release by a longhorn (cerambycid) beetle, which may be the small amounts of methane all organisms produce abiotically but the actual mechanism of this consistent methanogenesis remains intriguingly unknown.

**Originality-Significance Statement:** The research presented here goes beyond the long-established tracks on the symbiotic basis of insect methanogenesis and shows that there are areas still scarcely covered so far, with great exploratory potential.

## Introduction

The occurrence of symbiosis between arthropods and the microorganisms inhabiting their gastrointestinal tracts is a phenomenon widely studied in the case of termites and much less understood in other groups. Generally, in this type of relationship microorganisms participate in breaking down poorly digestible food and provide their hosts with nutrients. In return, symbiotic microorganisms are exposed to favourable, frequently anoxic environments (Brune, 1998) in which they can acquire energy-providing substrates (Nardi et al., 2002). As wood is generally rich in poorly digestible polymers (cellulose, hemicellulose, lignin) and low in nutrients (Haack and Slansky, 1987), wood-feeders are highly dependent on a plethora of symbiotic relationships with microorganisms (Price, 1984; Haack and Slansky, 1987).

The cellulose digestion process is best known for termites and is associated with a great number of distinct physiological groups of microorganisms, such as acetogenic non-spirochetal and spirochetal bacteria and methanogenic archaea. The latter group is of a special interest. The role of methanogens in the digestion process in wood-feeding termites is related to the presence of gut protists (Purdy 2007 and references therein), although a direct role in digestion is unclear. The specific spatial distribution of methanogens inside insect guts implies that their presence there is directly linked to the effective digestion of plant matter. Methanogens almost certainly act as hydrogen sinks and in some cases maintain the very low H_2_ partial pressures that are necessary for completion of organic matter degradation in the insect guts (Breznak & Brune, 1994, Brauman et al., 2000, Brune 2019). The by-product, and thus one possible diagnostic feature of intestinal methanogens presence, is methane emitting by their hosts (Breznak and Brune, 1994; Breznak, 2000; Brauman et al., 2000; Friedrich et al., 2001; Brune 2019). This phenomenon attracts much attention because methane is a potent greenhouse gas. Understanding of biological mechanisms of methane production in nature is important to create the mechanistic models, and to predict the scenarios of its future emissions (Conrad, 2020).

In 1994 Hackstein and Stumm examined 35 arthropod taxa and found that methanogenesis is limited to four groups of terrestrial arthropods: Diplopoda, Isoptera, Cetoniidae and Blattidae. This review included only one species of the strictly and exclusively wood-feeding longhorned beetles, *Hylotrupes bajulus* (Coleoptera: Cerambycidae). The exclusive wood-feeding within this group of insects makes them the very promising organisms for the studies on symbiotic methanogenesis but the tested beetles showed no methane release. However, this species is known to rely on proteins in food and to assimilate only about 35% of cellulose and 20-30% of hemicelluloses (Seifert, 1962, after Dominik & Starzyk, 2004). Unlike other cerambycid beetles, *H. bajulus* does not have symbiotic intestinal yeasts (e.g. Nardi et al., 2002), which implies that the mechanism of energy assimilation in this species is independent of complicated microsymbiotic relationships. Since the Hackstein and Stumm’s work, many groups of soil invertebrates were found non-methanogenic (Šustr and Šimek, 2009), while it is noteworthy that the list of methanogenic groups of arthropods has not been extended since then. Specifically, no other cerambycid beetle have been tested in this regard, according to our knowledge.

The symbiotic methanogens are either hydrogenotrophic, i.e., they reduce CO_2_ to methane by H_2_, or methylotrophic, those reducing the compounds containing a methyl group (Hackstein, 2010). So far, methanogenic archaea have been examined with molecular techniques in different termites (Ohkuma & Kudo, 1998, Ohkuma et al., 1999, Shinzato et al., 1999, Brauman et al., 2001, Friedrich et al., 2001, Donovan et al., 2004, Paul et al. 2012, Shi et al. 2015), cockroaches *Eublaberus posticus* (Cruden & Markovetz, 1987, after Sprenger et al., 2000), and *Periplaneta americana* (Sprenger et al., 2000), rose chafers *Pachnoda ephippiata* (Egert et al., 2003) and *Melolontha melolontha* (Egert et al. 2005), and millipedes (Šustr et al., 2014, Horváthová et al. 2021). The intestinal methanogenic symbionts of arthropods are derived from lineages representing four orders (Methanobacteriales, Methanosarcinales, Methanomicrobiales and Methanomassiliicoccales), out of eight euryarchaeotal orders of methanogens identified so far (Brune, 2019). They are found in different proportions in the gastrointestinal tracts of different arthropod groups (Brune, 2019), and among species representing one group, e.g., termites (Shi et al. 2015).

In this study, we explored whether the larvae of a cerambycid beetle *Stictoleptura rubra* L. are engaged in symbiotic relationship with methanogens. This species is known to live in symbiosis with the intestinal yeast *Candida tenuis* (Buchner, 1965, Jones et al., 1999). Its larvae live in dead coniferous trees and digest xylem, the least nutritious tissue of a stem. Previous studies show that the life history of this species is apparently limited by the nutritional quality of food, as showed in stoichiometric (Filipiak & Weiner, 2014, 2017; Filipiak, 2018) and bioenergetics and life history studies (Walczyńska 2007, 2008, 2010, 2012). Also, there is evidence that supplementing the larval diet with nitrogen resulted in a decreased metabolic rate, which suggests that it has some metabolically costly adaptations to overcome the nutritional deficiency of wood as a food type (Walczyńska 2009). Harnessing gut microbes to facilitate digestion of wood, similar to that employed by other methane-producing wood-feeding arthropods, may be a promising candidate for such an adaptation.

We used gas chromatography to detect any methane release and molecular techniques to directly search for the presence of intestinal archaea in the larvae of *Stictoleptura rubra.* Additionally, we used larvae of specimens of an unidentified buprestid beetle for comparison in gas chromatography.

## Materials and Methods

### Studied species

*Stictoleptura* (formerly, *Corymbia*) *rubra* (Linnaeus, 1758) is a relatively large cerambycid beetle species, the larvae of which feed on the most inner part of the stem of decayed coniferous trees, mostly pine tree (Dominik & Starzyk, 2004). This species grows slowly and develops for a relatively long time; the whole lifecycle lasts for 3-4 years (Walczyńska et al. 2010, Walczyńska 2012). This fact, accompanied by the study which showed its low assimilation efficiency index (Walczyńska 2007, 2008), points to the apparent specific adaptations of this species to the poor nutritional quality of its food source. For the study, larvae were obtained from decayed pine logs collected from the Niepołomice Forest near Krakow, Poland (50.04 N, 20.35 E). Larvae removed from logs were used alive for gas chromatography and frozen at −80°C prior to molecular analysis.

### Methane release – gas chromatography

Two measurement trials were conducted. In the first trial, larvae were maintained under laboratory conditions at constant temperature of 17° C for a few months, provided with wood originated from the stumps of their origin. In the second trial, larvae of *S. rubra* and, additionally, larvae of a wood-feeding buprestid beetle were collected in the field and examined on the same day. In both trials, weighed larvae were placed separately in test tubes with blotting paper moisturised with distilled water to maintain humidity during the experiment. All the test tubes were put into a glass container of 0.62L. In a second trial, larvae were additionally provided with their natural food in the form of pinewood cubes, about 5.5 g. Additionally, another glass container with similar amount of pinewood cubes only was tested at the same time to account for the possible effect of methane release by decaying wood. The second trial was subdivided in two, one initiated just after completing the other. As we aimed at finding any evidence of methane release, we analysed all available larvae exemplifying a wide range of sizes. Data on larvae weight, the number of individuals per container, and the duration of analyses are provided in Table 1A.

**Table 1.** Chomatographic analysis. **A.** Data on the individuals investigated in the gas chromatography experiment and experiments’ time duration. **B.** The rate of methane production by *S. rubra* (Cerambycidae) and buprestid beetle (Buprestidae) larvae.

| trial No | species | No of individuals | weight [g]<br>(mean ± SD [total]) | trial duration [hr] |
| --- | --- | --- | --- | --- |
| 1 | *S. rubra* | 16 | 0.20 ± 0.15 [3.24] | 20 |
| 2 | *S. rubra* | 10 | 0.18 ± 0.13 [1.82] | 41 / 79* |
| | buprestid beetle | 3 | 1.16 ± 0.14 [3.49] | 41 / 79* |

\* - duration of trial 2.1 / 2.2

**B.**
| trial No | sample type | methane concentration difference |  |
| --- | --- | --- | --- |
| | | ppb ± error* | nmol / hr×g ± error* |
| 1 | *S. rubra* | 126 ± 12 | 0.054 ± 0.005 |
| 2 2.1 | *S. rubra* | 83 ± 32 | 0.031 ± 0.012 |
| | buprestid beetle | -6 ± 47 | - |
| | pinewood | -13 ± 16 | - |
| 2.2 | *S. rubra* | 90 ± 8 | 0.017 ± 0.001 |
| | buprestid beetle | 25 ± 40 | - |
| | pinewood | 19 ± 15 | - |

\* - error was estimated as SD from mean value for all possible combinations of pairs of measurements

At the beginning and the end of each trial, two air samples per container were taken using a 50 ml gastight syringe and were injected in the Agilent 6890N gas chromatograph with a 1.5m Porapak Q packed column (Nęcki et al., 2003) equipped with a flame ionisation detector (FID) to detect methane. Prior to the experiment, syringes were purified and filled with synthetic air to avoid contamination of samples. Although only 5ml of gas was injected during a single analysis, approximately 40ml of sample gas was used to flush all the tubes and connections of the inlet system. The mole fraction of methane was calculated using a reference standard analysis performed every five sample analyses and expressed in a World Meteorological Organisation CH4 mole fraction scale (WMO, 2001). The accuracy associated with methane chromatographic analyses was <2 ppb; however, reproducibility of the whole analytical method can be assumed as 10 ppb. The quantity of released methane was determined as a difference in its mole fractions after 20, 41 or 79 hr of keeping larvae in an airtight container, in darkness at room temperature. Data were analysed using a bootstrap method (Tong et al., 2012) in R environment in the ‘ClusterBootstrap’ package (Deen and de Rooij, 2020), with an assumed twenty times increase in sample size. The differences in methane release by *S. rubra* larvae between trials 1 and 2, as well as among different type of examined samples (*S. rubra* larvae, buprestid larvae, pinewood and blank) in trial 2, were tested using a t-Weltch test with an option of different degrees of freedom.

### Presence of gut Archaea – 16S rRNA gene sequencing and T-RFLP analysis

The weight range of larvae was 0.01-0.71 g. The larvae were kept frozen until their guts were removed. DNA was extracted from whole beetle guts (in batches of 6-20) by using the hydroxyapatite spin-column method (Purdy, 2005) and purified with PEG 6000. The euryarchaeal 16S rRNA genes were amplified in nested PCR and analysed by T-RFLP as in Donovan et al. (2004). For T-RFLP analysis the labeled primer 1100R-FAM was used. The fluorescently labeled PCR products were digested by using the 4-bp cutter TaqI (Promega, Southampton, United Kingdom) as described by the manufacturers. About 2 ng of the restricted PCR product was analysed on an ABI 377 automatic sequencer (Sequencing Facility, Natural History Museum, London, United Kingdom). For the clone library, the purified PCR product from the secondary, unlabeled PCR was cloned into pGEM-T Easy (Promega) and positive clones were checked by amplification with the vector-based primers M13f and M13R.

## Results

### Methane release

Larvae of *S. rubra* released methane at the average rate of 0.03 nmol/g × hr (Table 1B). After completion of trial 2.2, two larvae of *S. rubra* and one larva of a buprestid beetle were dead. In two cases, one air sample was omitted from analyses because of error in sample dosing (trial 1) or because of apparent difference between one of the measurements and other three (> 100 ppb; buprestid larvae, trial 2.1). Methanogenesis of *S. rubra* larvae did not differ between trials 1 and 2 (p = 0.1; Table 2). In trial 2, methane production from the *S. rubra* larvae was significantly greater than that from the controls (p <0.001) and pinewood (p = 0.046) and it tended to be greater than that from buprestid larvae (p = 0.056; Fig. 1; Table 2). Neither buprestid larvae nor pinewood differed significantly from controls (Table 2).

**Fig. 1.**
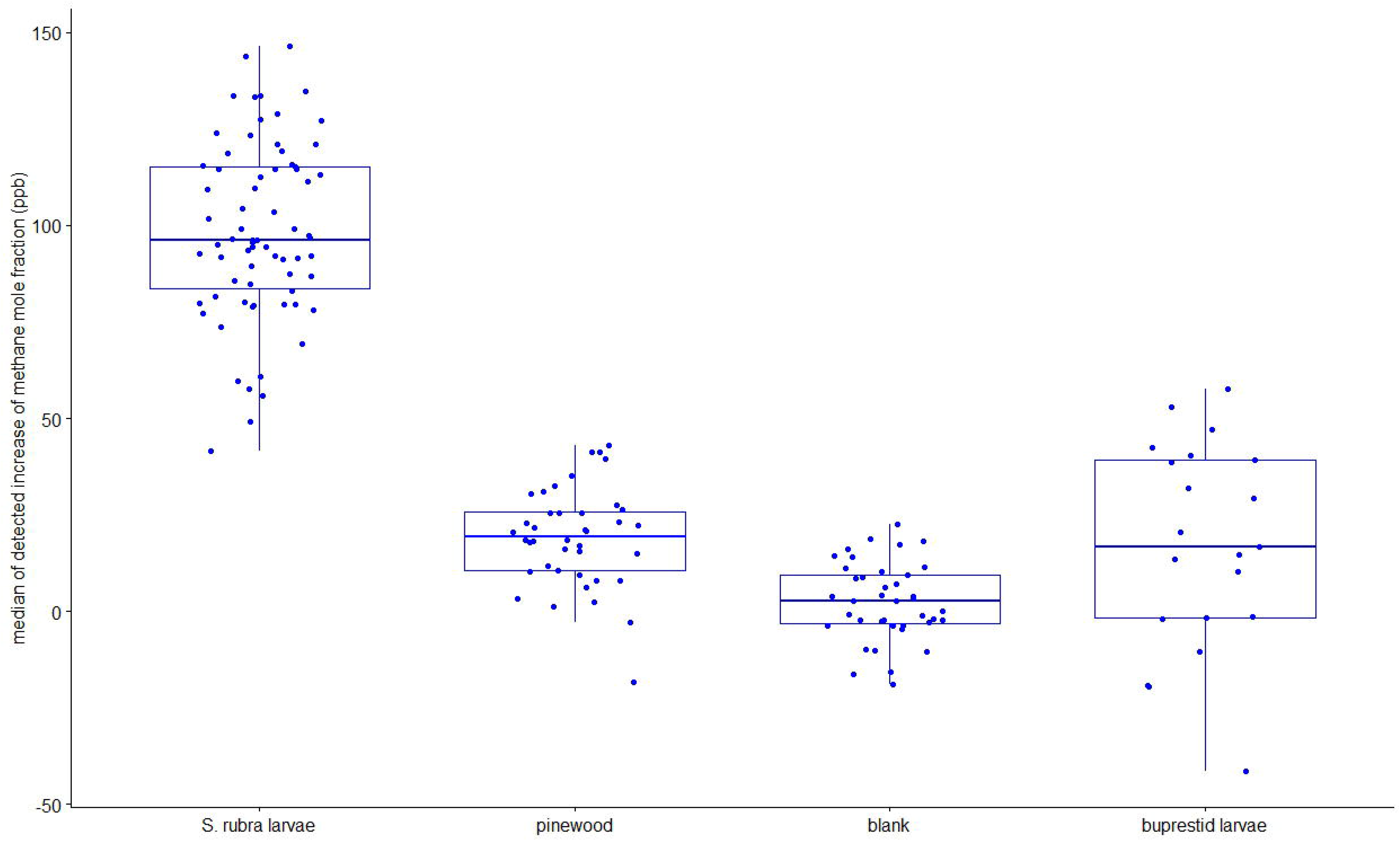
The box-plot graph of methane mole fraction enhancement (ppb) noticed for different types of samples analysed and bootstrapped. horizontal line = median, box = quartiles 2 and 3, vertical lines = quartiles 1 and 4, dots = individual values

**Table 2.** Statistical results of t-Welch test after bootstrap analysis (bootstrapped 10000 ×). Sample 1 and 2 denote the two compared sample types; N1 and N2 denote the original sample size.

| sample 1 | sample 2 | N1 | N2 | df | t | p-val |
| --- | --- | --- | --- | --- | --- | --- |
| blank | *S. rubra* | 4 | 7 | 6.6 | 6.7 | 0.00011 |
| pinewood | *S. rubra* | 4 | 7 | 4.5 | 3.166 | 0.046 |
| buprestid larvae | *S. rubra* | 2 | 7 | 2.03 | 3.88 | 0.056 |
| buprestid larvae | pinewood | 2 | 4 | 3.2 | 0.44 | 0.66 |
| buprestid larvae | blank | 2 | 4 | 1.05 | 0.055 | 0.87 |
| pinewood | blank | 4 | 4 | 3.9 | 1.58 | 0.58 |
| *S. rubra* trial 1 | *S. rubra* trial 2 | 2 | 5 | 3.1 | -6.37 | 0.1 |

### Presence of Archaea

We detected clones very closely related to the obligately halophilic *Halorubrum saccharovorum* (Halobacteriales, Euryarchaeota), with their phylogenetic position provided in Fig. 2.

**Fig. 2.**
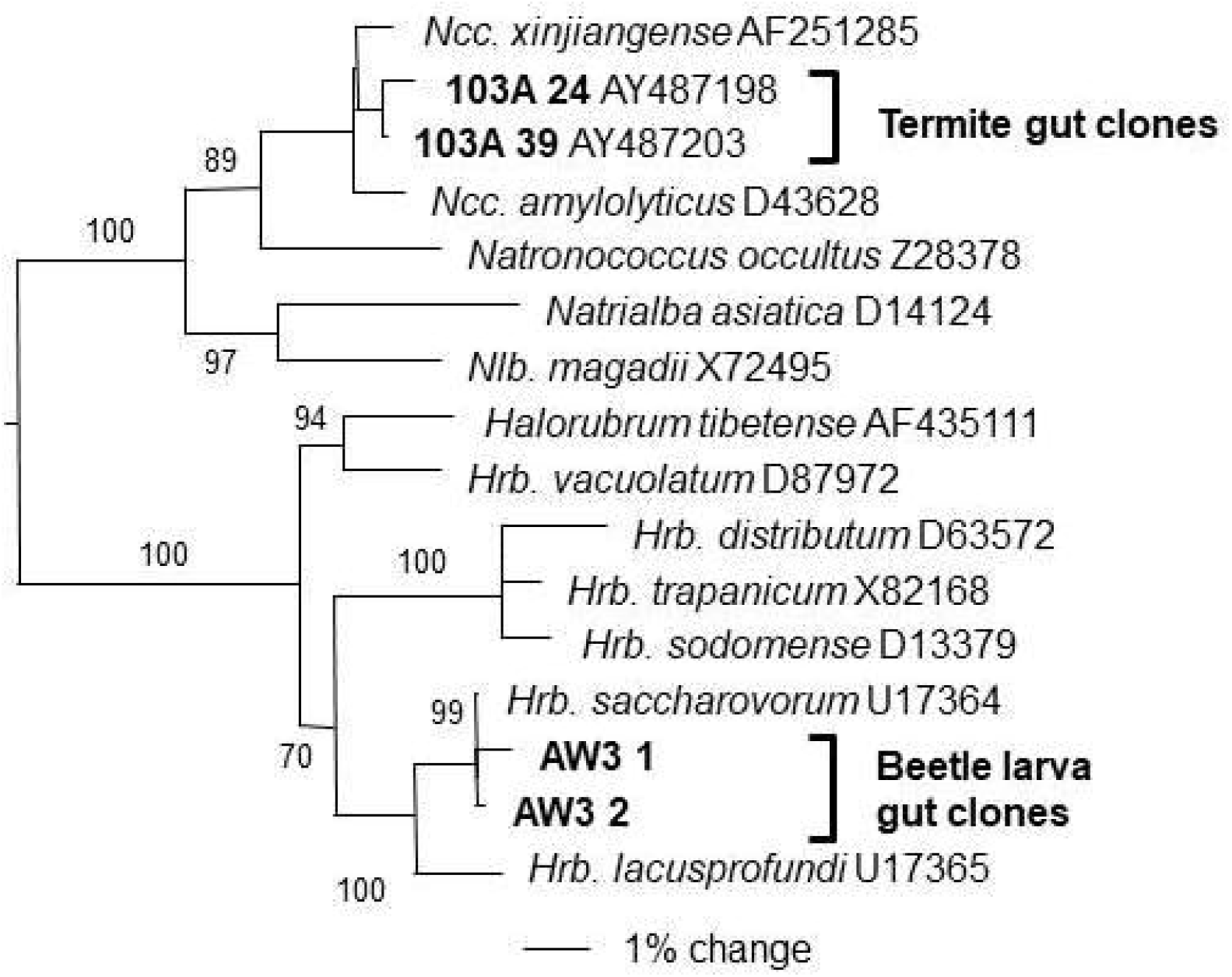
Neighbour joining tree showing the phylogenetic relationships between clones from the gut of *Stictoleptura rubra* larvae, a soil termite gut and members of the halophilic Halobacteriales. Numbers on nodes are bootstrap values from 100 iterations.

## Discussion

We report the first case of methane release in the longhorned beetle *Stictoleptura rubra* (Cerambycidae) larvae. Though the amounts of methane released are small, a few orders of magnitude smaller than in other methanogenic arthropods, they are methodologically significant, being several times larger than the methane detection limit for methane mole fraction changes at the ambient laboratory level. Additionally, the comparison with another wood-feeding beetle which appeared to be non-methanogenic supports the claim that methanogenesis in *S. rubra* represents not simply physiological background but it may rather be the result of a symbiotic relationship with methanogens. Biological causes of variation in methanogenesis rate obtained in trials 1 and 2 might have resulted from the presence of wood in the trial 2, which could have absorbed some methane, decreasing the observed release rate. Variation in data obtained in trials 2.1 and 2.2 may result from differences in their duration linked with an increase in CO_2_ concentration in the trial 2.2 in comparison with the trial 2.1 (data not shown) or from processes connected with decay of dead larvae. However, methane production from the cerambycid beetle larvae was significantly greater than that from the controls.

The results of molecular search for archaeal genes in the gut of *S. rubra* larvae provide the intriguing results. The *Halobacteriales* are clustered between class I class II methanogens (Paul et al. 2012) and clones from this order have been identified previously from the gut of *Cubitermes fungifaber* termite (Donovan et al. 2004), but their relationship with the hosts (whether symbiotic or other) is not certain and cultured members of the halobacteriales are all non-methanogenic. This archaeal group is suggested to have secondarily lost the methanogenic metabolic pathway (Bapteste et al. 2005). This result leaves open the question where did the methane detected come from if there are no methanogens present? While highly speculative, it could be explained if the *Halorubrum*-like sequences detected in the longhorn beetle larva gut are from an unknown methanogen. The discovery of the *Methanomassiliicoccales* within the *Thermoplasmalates* clade (Dridi et al. 2006, Wright, et al. 2011) indicates that it cannot be assumed that all possible methanogens have been detected.

The role of *S. rubra* as a natural methane-emitter seems negligible, but its relationships with intestinal microorganisms are worth being investigated more closely. Providing this species does produce methane through symbiotic organisms, the low levels of methane emissions may be related to it being a temperate species because it is known that temperature affects the methanogenic pathways in a way that hydrogenotrophic methanogenesis is favoured at higher temperatures (Conrad, 2020). Therefore, tropical arthropods may emit more methane than temperate ones. Also, methanogenesis in non-termite arthropods seems to be flexible and dependent on different factors. In millipeds for example, it was found to change with food type and body size, among other factors (Šustr et al. 2020).

This short study was conducted because of pure curiosity and it does not reflect the mainstream of our research. We expected either a negative result or a positive one, being in line with the current state of knowledge. To our surprise, we obtained results that may form the basis for a new research pathway within methanogenesis resulting from the symbiosis of wood-boring insects with gut microbes.

## Acknowledgements

The authors thank Paul Eggleton for helpful comments and the Sequencing Facility at the Natural History Museum in London for sequencing. The molecular work was performed at the University of Reading, Department of Animal and Microbial Sciences, funded by a grant from the British-Polish Young Scientists Programme to AW. The chromatographic part of research have been developed with the use of equipment financed from the funds of the “Excellence Initiative – Research University” program at AGH University of Science and Technology. AW thanks Professor January Weiner for his idea to conduct this study. The authors declare no conflict of interest.

## Notes

### Competing Interest Statement

The authors have declared no competing interest.

